# Highly resolved genomes as a tool for studying speciation history of two closely related louse lineages with different host specificities

**DOI:** 10.1101/2023.11.03.565481

**Authors:** Jana Martinů, Hassan Tarabai, Jan Štefka, Václav Hypša

## Abstract

Sucking lice of the suborder Anoplura are permanent ectoparasites with specific lifestyle and highly derived features. Currently, genomic data are only available for a single species, the human louse *Pediculus humanus*. In this study we present genomes of two distinct lineages, with different host spectra, of a rodent louse *Polyplax serrata*. Genomes of these ecologically different lineages are closely similar in gene content, display a high level of synteny, but they also differ by a few duplications/translocations and single inversion. Compared to *P. humanus*, the two *P. serrata* genomes are noticeably larger (139 Mbp vs. 111 Mbp) and encode a higher number of genes. Similar to *P. humanus*, they are significantly reduced in sensory-related categories such as vision and olfaction. Utilizing a genome-wide set of genes, we perform phylogenetic reconstruction and evolutionary dating of the *P. serrata* lineages. Obtained estimates reveal their relatively deep origin (approx. 6.5 Mya), comparable to the time of split between the human and chimpanzee lice *Pediculus humanus* and *P. schaeffi*. This dating supports the view that the *P. serrata* lineages are likely to represent two cryptic species with different host spectra. Historical demographies of the two lineages show glaciation-related population size (Ne) reduction, but recent restoration of Ne was seen only in the less host specific lineage. Together with the louse genomes, we analyze genomes of their bacterial symbiont *Legionella polyplacis*, and evaluate their potential complementarity in synthesis of amino acids and B vitamins. We show that both systems, *Polyplax*/*Legionella* and *Pediculus*/*Riesia*, display almost identical patterns, with symbionts involved in synthesis of B vitamins but not amino acids.

## Introduction

Among several thousands of insect genome records currently available in NCBI, only six represent permanent ectoparasites of the order Phthiraptera, and just two the suborder of sucking lice Anoplura, both represeting a single species *Pediculus humanus*. Of these genomes three have been analysed and published in the form of a genomic comparative study. One of them, *Pediculus humanus* (Kirkness et al., 2010), represents sucking lice, and the other two, *Columbicola columbae* and *Brueelia nebulosa*, are chewing lice of the family Philopteridae (Baldwin-Brown et al., 2021; Sweet, Browne, Hernandez, Johnson, & Cameron, 2023). In this study, we reconstruct genomes of two phylogenetically distinct lineages of the sucking louse *Polyplax serrata*, providing genomic data for a second anopluran species and allowing for the comparison between different sucking lice genera. Such comparison is important to distinguish which features of *P. humanus* stressed in the previous comparative analyses of phthirapteran genomes (Baldwin-Brown et al., 2021; Kirkness et al., 2010; Sweet et al., 2023) represent common anopluran characteristics and which are unique to this species (e.g., highly reduced genome size, loss of sensory genes etc.).

In addition to this general significance, the genomes of the two *P. serrata* lineages provide an important background for evolutionary studies of this louse species. The lice classified as *P. serrata* were shown in series of studies to form a complex assembly of populations with different distributions and ecologies (Martinu, Hypsa, & Stefka, 2018; Martinu et al., 2015; Martinu, Stea, Poosakkannu, & Hypsa, 2020). The most conspicuous difference across this assemblage is the degree of host specificity in two different lineages, namely the “non-specific lineage” (N lineage) parasitizing two host species (*Apodemus sylvaticus* and *A. flavicollis*) and the “specific lineage” (S lineage) restricted only to *A. flavicollis*. In fact, the phylogenetic and genealogical paterns obtained in the previous analyses indicate that these lineages are likely to represent two different but morphologically undiscernible species.

Finally, like all other sucking lice, *P. serrata* maintains obligate symbiosis with a bacterial symbiont *Legionella polyplacis* (Rihova, Novakova, Husnik, & Hypsa, 2017). Such mutualistic associations with bacteria are known from various ecological types of insects. Genomes of these symbionts typically undergo dramatic changes, mostly losses of genes, and their metabolic capacities evolve to reflect the host’s lifestyle and source of the diet (Kinjo et al., 2022). In blood-feeding insects, complete genomes of the host and the symbiont have so far been available only for tsetse flies of the genus *Glossina* and their symbiont *Wigglesworthia glossinidia* (Akman et al., 2002; Rio et al., 2012), and for *Pediculus humanus* with the symbiont *Riesia pediculicola* (Kirkness et al., 2010).

## Results and discussion

### Genome assemblies and annotation

Two high-quality genome assemblies were generated combining Oxford Nanopore (coverage 108x for S lineage and 42x for N lineage) and Illumina data. Genome assemblies of the two *P. serrata* lineages yielded similar main characteristics (Table 1). For the S lineage, the total assembly size was 138.66 Mbp with the largest scaffold covering 20.69 Mbp, and scaffold N50 of 10.50 Mbp. The genome encoded 14,045 predicted genes, 13,914 mRNA and 131 tRNA. For the N lineage, the assembly reached 138.54 Mbp, the largest scaffold covered 17.59 Mbp, and scaffold N50 was 13.3 Mbp. BUSCO genome completeness analysis found 98.7% (1000/1013) and 98.9% (1002/1013) complete BUSCOs of *P. serrata* S and N lineages, respectively (Supplementary Table S1, Datasheet S1). Gene prediction of N lineage revealed 15,132 genes, 14,991 mRNA and 141 tRNA. The repeat sequences constituted 4.93 Mbp (3.56%) of the genome size in S lineage, and 4.84 Mbp (3.49%) in the N lineage. In both the S and N lineages, simple repeats constituted 49% of all identified repeat elements, while interspersed repeats accounted for 40% in each lineage. A slightly higher difference was observed for the content of long interspersed nuclear elements (LINEs), which constituted 0.52% of total genomic length in S lineage compared to 0.39% in N lineage (see Supplementary Table S1, Datasheet S2 for detailed breakdown of the identified repeat elements).

**Table 1.** Overview of the genome characteristics of the analyzed Phthiraptera.

| Genome characteristics | *Polyplax serrata* S | *Polyplax serrata* N | *Pediculus humanus* | *Brueelia nebulosa* | *Columbicola columbae* |
| --- | --- | --- | --- | --- | --- |
| Assembly length | 138,661,288 | 138,547,654 | 110,770,411 | 113,965,818 | 207,887,661 |
| Num Scaffolds | 89 | 70 | 1,873 | 1,684 | 384 |
| Percent GC | 36.90% | 36.92 | 26.87% | 37.84% | 36.11% |
| N50 | 10,507,470 | 13,306,208 | 497,057 | 636,874 | 17,673,050 |
| Num Genes | 14,045 | 15,132 | 12,130 | 15,901 | 25,246 |
| Num Proteins | 13,914 | 14,991 | 12,004 | 15,814 | 25,113 |
| Num tRNA | 131 | 141 | 126 | 87 | 133 |
| Average gene length | 2,667.1 bp | 2,335.4 bp | 2,573.6 bp | 2,248.2 bp | 2,097.8 bp |
| Transcript CDS | 13,914 | 14,991 | 12,004 | 15,814 | 25,113 |
| CDS complete | 13,848 | 14,520 | 11,493 | 15,216 | 24,485 |
| Total exon no. | 76,836 | 78,485 | 71,562 | 83,006 | 109,157 |
| Total exon CDS | 76,836 | 78,485 | 71,562 | 83,006 | 109,157 |

In contrast to the high similarity of the two closely related S and N lineages, *P. serrata* differs considerably from *P. humanus*, the only other sucking louse for which genome assembly is available (Table 1). Generally, *P. serrata* genomes are larger, with a higher number of genes and a higher GC content. However, the differences between the two anopluran genera are relatively small when compared to the differences between the two chewing lice included in the study. While the genome of *B. nebulosa* is larger but comparable in size to the anopluran genomes, the genome of *C. columbae* is almost twofold larger.

### Comparative genomic analysis

The genomes of the two phylogenetically distant species, *P. serrata* and *P. humanus* share a high proportion of their protein contents. Over 93% of the annotated proteins from the PFAM and InterProScan databases were conserved across the three analyzed anopluran genomes (Figure 1, Supplementary Table S1, Datasheet S3 and S4). The analyses using several different databases provided similar results, indicating a high number of shared features and only few unique features (Figure1; Supplementary Table S1, Datasheets S5-S7), with exception of predicted signal peptides, showing lower number in *P. humanus* (n=702) compared to that of *P. serrata* S (n=1,049) and N (n=1,063) (Figure 1, Supplementary Table S1. Datasheet S7).

**Figure 1.**
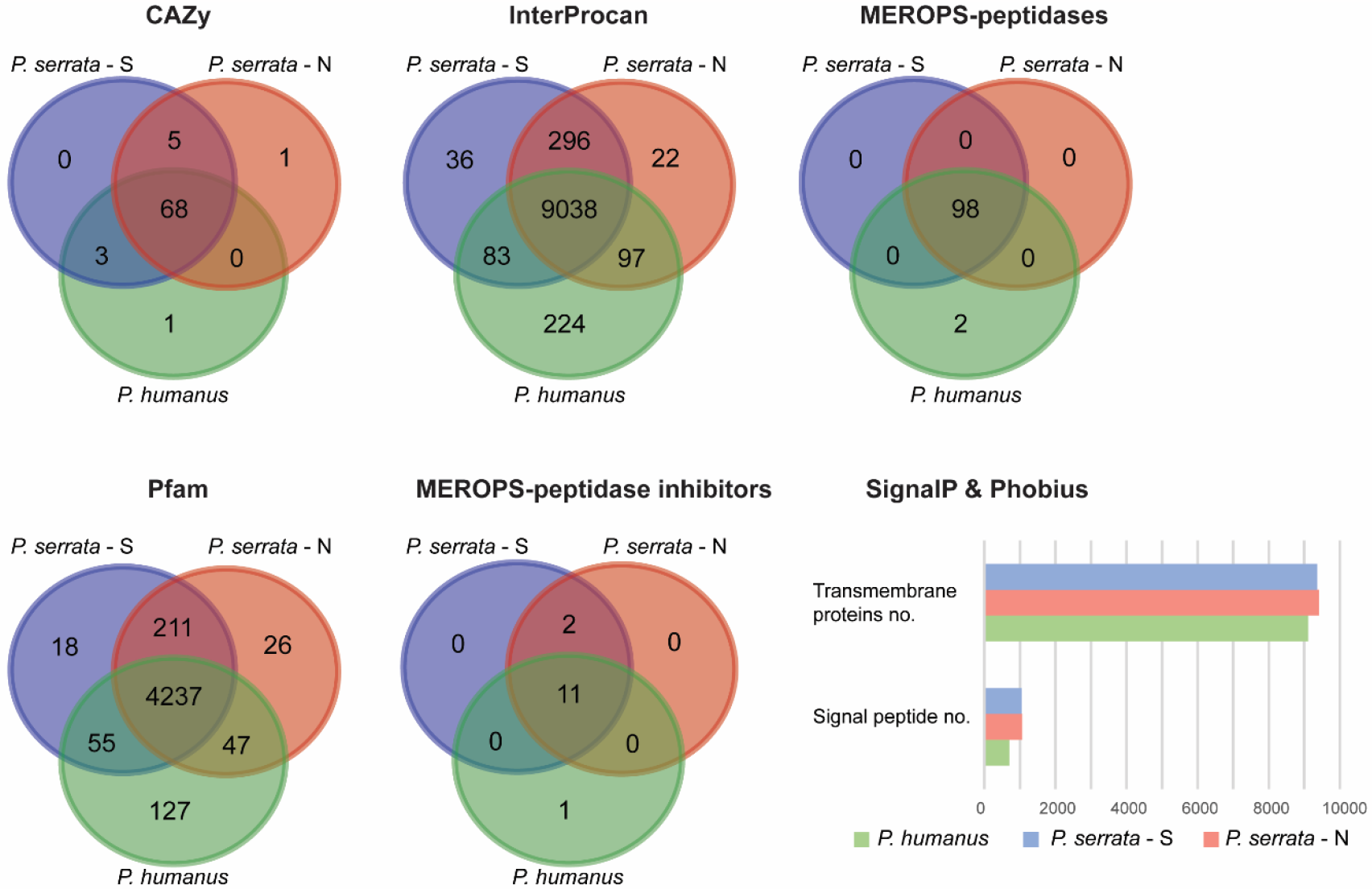
Comparison of genome contents of the three sucking lice (Anoplura). For the databases CAZy, InterProscan, MEROPS, Pfam, the numbers represent unique IDs identified in the genomes. For SignalP & Phobius the plot shows total numbers of the transmembrane proteins and signal peptides identified by the databases.

Comparison of the three anopluran genomes with the two available genomes of chewing lice and other blood-feeders is more complex (Supplementary figure 1; see Supplementary Table S1, Datasheet S8 for list and references of all compared genomes). In their analysis Baldwin-Brown et al. (2021) pointed out that chewing louse *Columbicola columbae* and sucking louse *P. humanus* possess reduced numbers of opsins (2 and 3, respectively). This is a much lower number than we detected in some blood-feeding dipterans (e.g., 20 in *Aedes aegypti*, 11 in *Glossina morsitans*), but comparable to blood-feeding heteropterans *Cimex lectularius* (2) and *Rhodnius prolixus* (5). In the *P. serrata* genomes presented here, the opsin genes (InterProScan IDs: IPR027430, IPR001760, and IPR001391) were entirely missing, reflecting the fact that in this species eyes are completely reduced.

An even stronger effect of the permanent parasitism on gene loss was observed in the repertoire of olfactory receptors (IPR004117). We obtained this annotation for only 18 genes in *B. nebulosa*, 21 in *C. columbicola*, 13 in *P. humanus*, and 15 in both *P. serrata* lineages, compared to much higher numbers in temporary insect parasites, specifically 38 in *Cimex lectularius*, 49 in *Glossina morsitans*, 66 in *Aedes aegypti*, and 153 in *Rhodnius prolixus*. We also detected a single gene associated with taste receptor activity (GO: 0008527) in each of the *P. serrata* lineages. In agreement with the Baldwin-Brown et al. (2021) report, we found two such genes in the *C. columbae* genome, but we did not detect this GO in *P*.*humanus*. However, when considering all annotations defined as “taste connected”, we found 16 genes in *B*.*nebulosa* and *C*.*columbicola*, 5 in *P*.*humanus*, 13 in *P*.*serrata* N, and 10 in *P*.*serrata* S. Apart from these differences in the repertoire of insect genes, we also observed various numbers of Rhabdovirus-related genes. Specifically, we identified 33 hits in *P*.*serrata* N lineage, compared to 10 hits in S lineage, 3 hits in *P. humanus* and single hit in *C. columbae* (Supplementary Table S1, Datasheet S4). This group of viruses is known to be broadly distributed across wide range of organisms, including plants, insects, and vertebrates (Ammar, Tsai, Whitfield, Redinbaugh, & Hogenhout, 2009). It was also demonstrated that in insects the rhabdoviruses can be vertically transmitted to progeny (Longdon et al., 2017). It is therefore difficult to hypothesize on a possible evolutionary or ecological significance of this finding in our data.

When visualizing genome dissimilarities of the five phthirapteran species by Principal Coordinate Analysis (PCoA), the plots well reflected their phylogenetic relationships and evolutionary distances. For the results obtained from the family-centered pfam database, the pattern was straightforward with the two *P*.*serrata* positioned as two closest points, and *B. nebulosa* as the genome most distant from all others (Figure 2A). In the InterProScan-based PcoA, the distant position of *B*.*nebulosa* lowers resolution between the anopluran genera (Figure 2B). However, after the removal of the *B*.*nebulosa* outlier, the analysis reveals close proximity of the two *P*.*serrata* lineages in comparison to significantly more distant *P. humanus* (Figure 2C).

**Figure 2.**
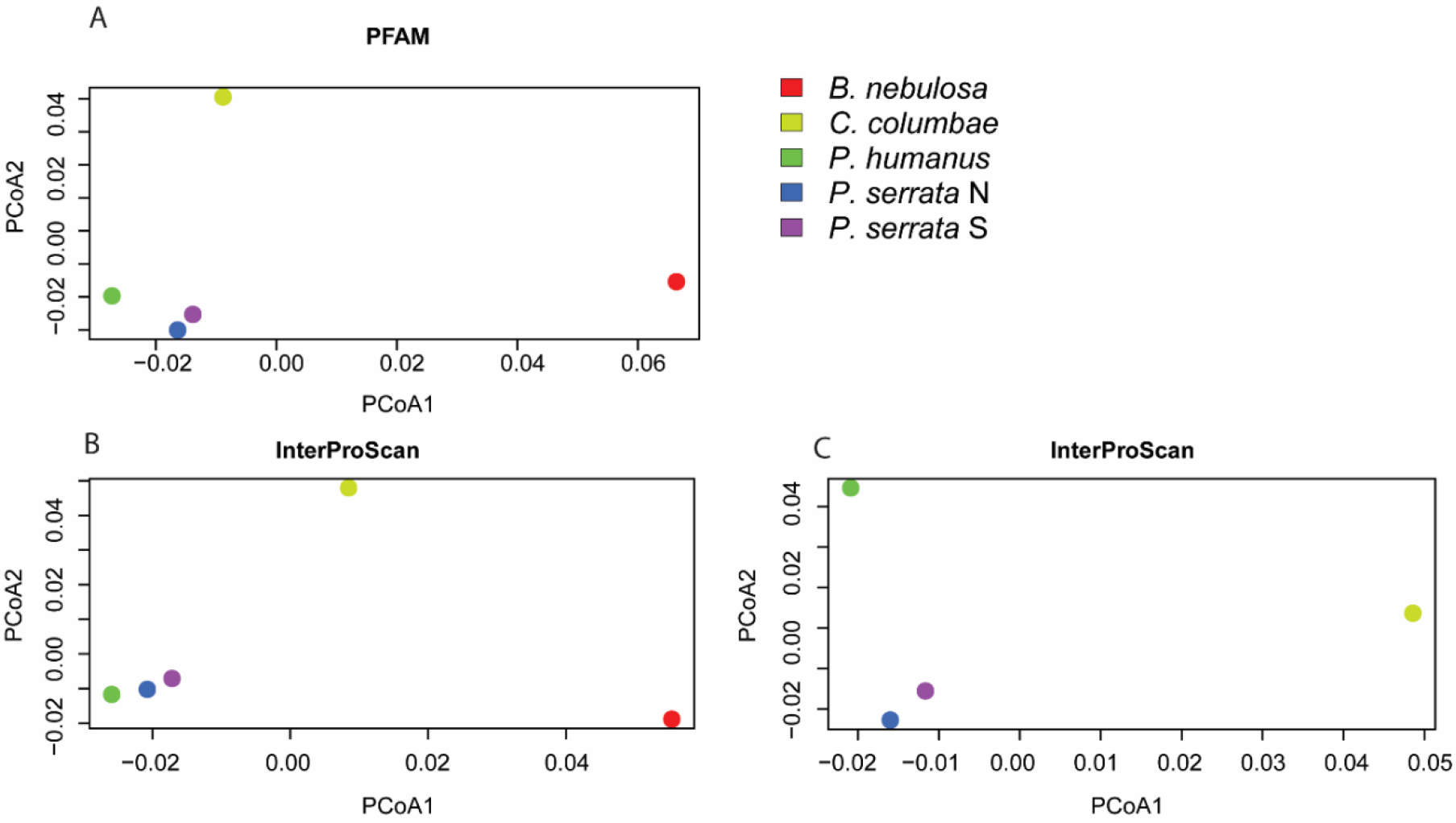
PcoA analysis of the five phthirapteran genomes. A – PCoA based on the results of the family-centered pfam database; B – PCoA based on the InterProScan database; C - PCoA based on the InterProScan database without outlier B. nebulosa.

### Synteny and chromosomes

Considering the genome size around 139 Mbp in the *P. serrata* S and N lineages (Table 1) and the number of chromosomes in *Polyplax* lice (Golub & Nokkala, 2004), we presume that the longest scaffolds in both *P. serrata* lineages (17-20 Mbp) likely represent almost complete chromosomes. High contiguity of the two assemblies allowed for comparing their synteny on a macroscale level. Contigs longer than 0.7 Mbp were chosen for collinearity analyses. Pair-wise comparison of the 18 longest contigs of S lineage (99.3% of the genome) and 21 of N lineage (98.7%) revealed a high degree of synteny with 78.14% of collinear genes (Figure 3A). Structural rearrangements occurred only between scaffolds PS6 in *Polyplax* S lineage and PN5, PN11, PN12 and PN18 of *Polyplax* N lineage (Figure 3B). More specifically, a 2.9 Mbp long fragment was translocated from PS6 to PN5 scaffold in inverted orientation, and several duplication/translocation events, involving short fragments of 2-11 genes, occurred between PS6 and the four *Polyplax* N contigs mentioned above. As high level of synteny is crucial for sexually reproducing species during recombination, its loss can have large impact on the success of mating and viability of the progeny (Feulner & De-Kayne, 2017; Simakov et al., 2020). Moreover, evidence from different study systems proposes chromosomal inversions as important drivers during speciation process with gene flow (McGaugh & Noor, 2012; Twyford & Friedman, 2015), which supports the idea that translocated and inverted parts of PS6 may serve as an intrinsic barrier to gene flow between *Polyplax* S and N despite the absence of morphological differences between the lineages and the sharing of one of their host species (Martinu et al., 2018; Stefka & Hypsa, 2008).

**Figure 3.**
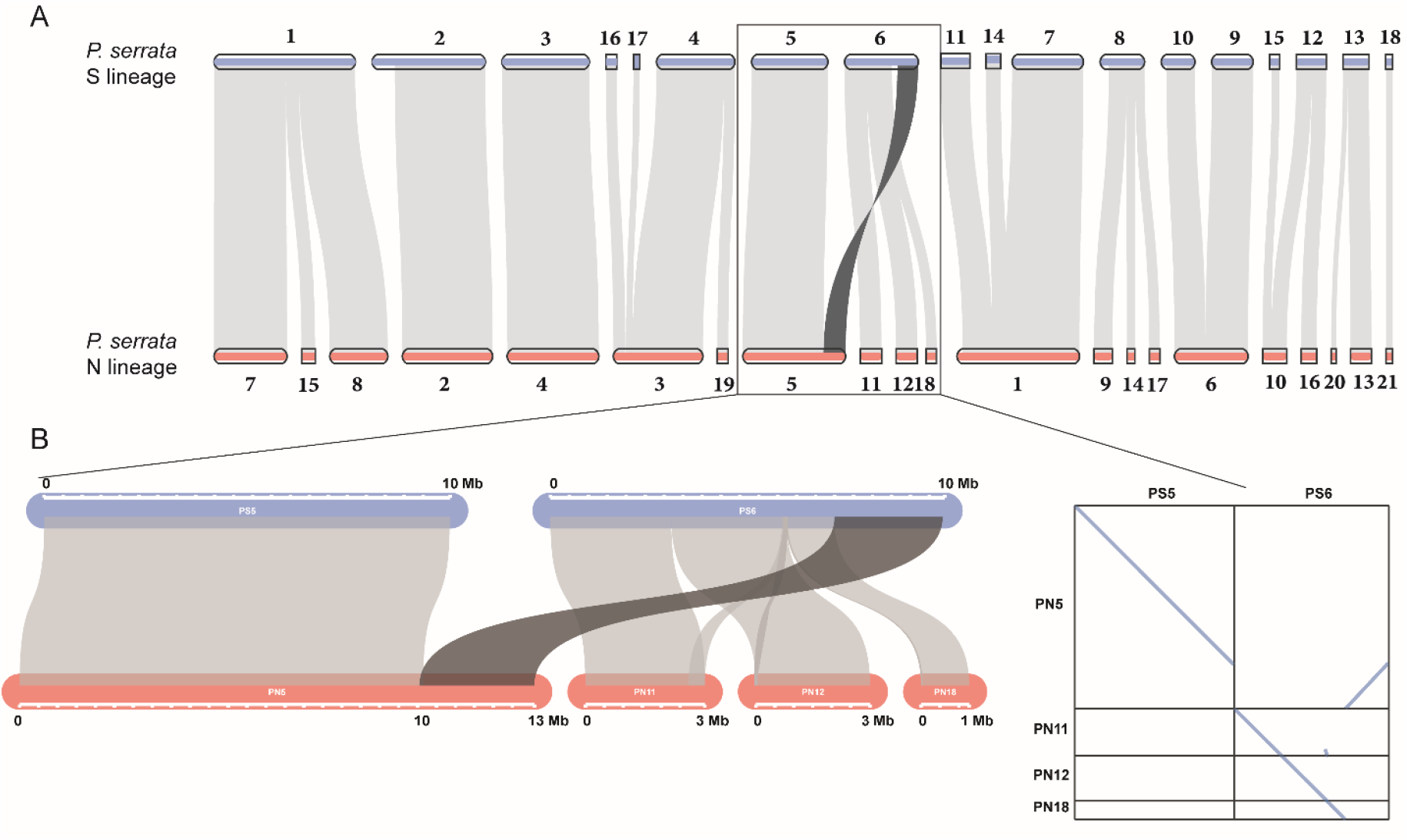
Dual synteny plot of longest scaffolds of P. serrata S lineage (blue) and N lineage (red). A) Syntenic regions identified by McScanX are highlighted by light grey, the inversion by dark grey. B) Detailed synteny and dot plots of structural variations between the scaffolds PS5-PS6 from S lineage and PN5-PN11-PN12-PN18 from N lineage visualized by SynVisio.

**Figure 4.**
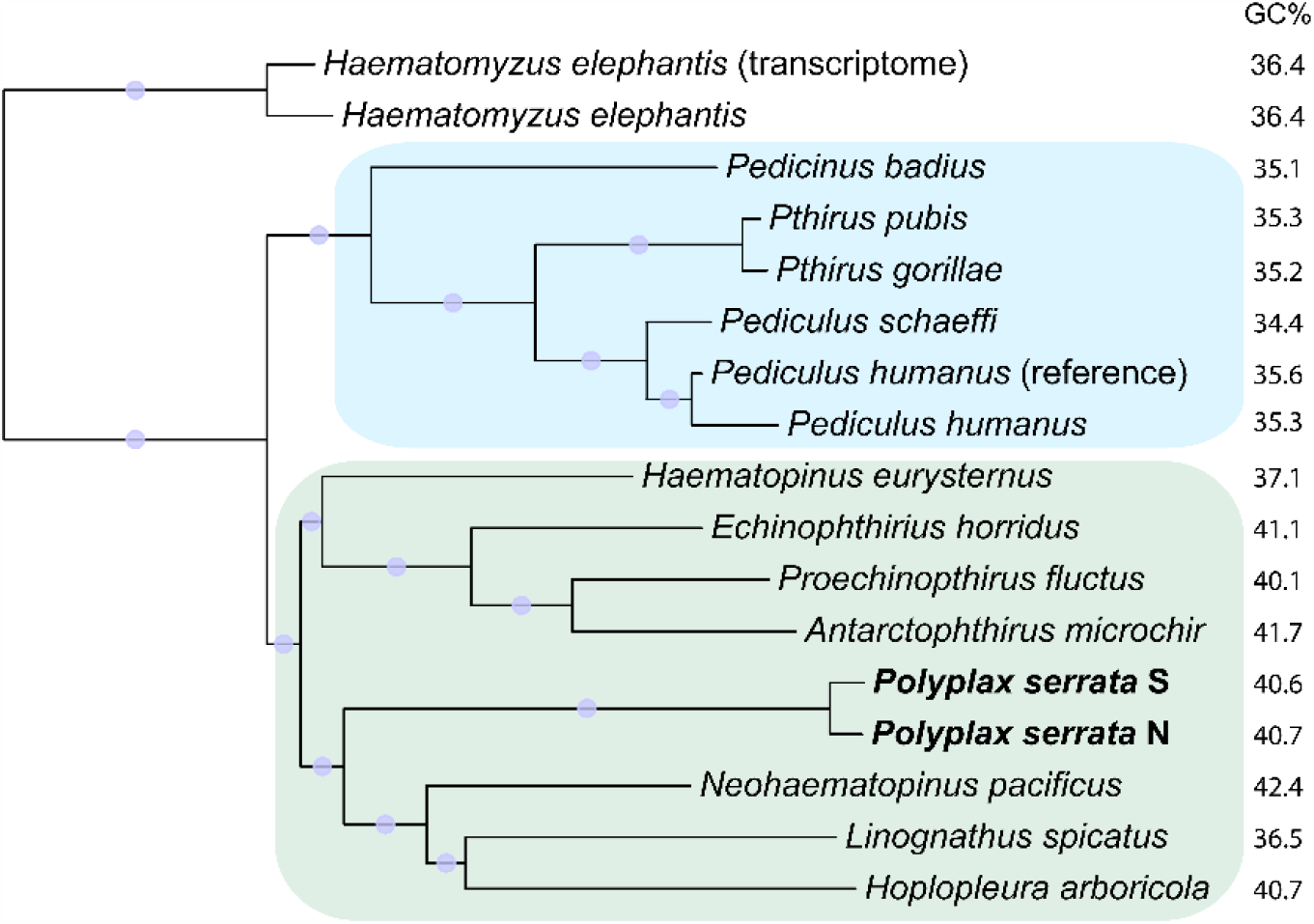
Phylogenetic tree derived by ML (iqtree2) from a matrix of 1049 orthologs (only 2nd codon positions; 536,051 sites). The new Polyplax genomes are printed in bold. GC% = GC content in full concatenated matrix (i.e., all codon positions). All bootstrap values reached 100 (indicated by the purple dots). The colored background designates the two branches with different ranges of GC content: blue = primate associated lice, green = lice associated with carnivores, ungulates, and rodents.

### Genome of Legionella polyplacis and host-symbiont complementarity

Both sucking lice, *P. serrata* and *P. humanus*, have previously been found to live in obligate symbiosis with bacteria *Legionella polyplacis* and *Riesia pediculicola*, respectively (Allen, Reed, Perotti, & Braig, 2007; Rihova et al., 2017). While our *P. serrata* hybrid Nanopore-Illumina assemblies contained only fragmented genomes of *L. polyplacis*, Spades assemblies of Illumina reads produced complete circular genomes (529,751 bp and 530,980 bp). The genomes were highly similar to the *L. polyplacis* samples reported previously (Martinu et al., 2020; Rihova et al., 2017). To compare the host-symbiont associations in *Polyplax* and *Pediculus*, we analyzed metabolic capacities in two categories usually considered in relation to the insect-bacteria symbiosis, B vitamins and amino acids. For both categories, the two anopluran genera show high similarity (table 2). The capacity for amino acids synthesis is in both lice determined almost strictly by the host genomes, while in the symbionts the pathways are largely deteriorated. The only difference between the two systems consists in the enzyme allowing conversion between serine and glycine, present in the *Riesia* genome of *Pediculus*. This amino acid patern corresponds to the general view that, in contrast to the insects feeding on plant saps, the blood feeding groups do not depend on their symbionts for amino acids. In contrast, at least three B vitamins (riboflavin, biotin, and folate) are provided by the symbionts. In both louse systems, the bacterial folate pathway lacks only one of the ten required enzymes (according to the KEGG module M00126). This high degree of conservation indicates that the pathway is functional and the missing reaction is fulfilled by an unknown gene/enzyme (Říhová, Gupta, Darby, Nováková, & Hypša). Comparison with the assembled *P. serrata* genome shows that one possible candidate is the louse alkaline phosphatase. The simple lipoic acid pathway is coded independently by both lice and the symbionts. A clear difference between the *Polyplax* and *Pediculus* systems represents the pantothenate pathway. While none of the symbionts code for this vitamin in their genomes, *Riesia* has been reported to carry a plasmid with the pantothenate genes (Boyd, Allen, de Crécy-Lagard, & Reed, 2014).

**Table 2.**
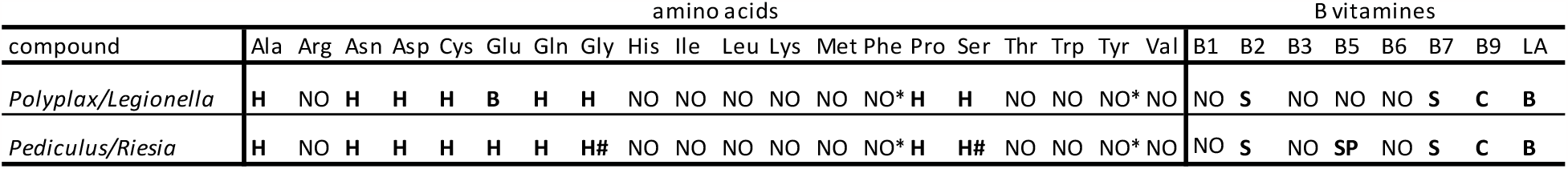
Metabolic capacities of the host/symbiont systems for amino acids and B vitamins. H = full pathway encoded by the host (H# = serine/glycine conversion capacity in Riesia); NO = the pathway is incomplete in the whole host/symbiont system (NO* = enzyme for Phe/Tyr conversion present in Polyplax); S = the pathway is coded by the symbiont; C = possible complementarity (potential role of the host’s alkaline phosphatase); B = full pathway is present in both counterparts; SP = pathway encoded on symbiont’s plasmid. The complete potentially functional pathways are printed in bold.

### Phylogeny and dating

Using ML based analysis we placed *P. serrata* lineages into the phylogeny of Anopluran taxa for which genomic data are available. The analysis produced a tree with strong bootstrap supports (Figure 3). Its topology agrees with the arrangement of anopluran species in the tree published by de Moya et al. (2021). The first dichotomy lies between the primate-associated lice (*Pedicinus, Pediculus, Phthirus*) and the remaining taxa. In the later clade, the two *Polyplax* lineages branched as a sister taxon to the *Hoplopleura*+*Linognathus*+*Neohaematopinus* cluster. This position of *P. serrata* is in conflict with the phylogeny previously published (Light, Smith, Allen, Durden, & Reed, 2010) which placed *P. serrata* as a sister group of the primate-associated genera. Since the tree we present here is based on a large amount of data, and supported by high bootstrap values, we consider this topology a reliable representation of the *P. serrata* position within Anoplura. Moreover, this topology better reflects differences in the GC content of the analyzed genomes. Since complete genomes are not available for most of the included Anoplura taxa, we used GC content of the selected set of genes as a proxy. The comparison shows that the striking difference between the *P. humanus* and *P. serrata* genomes in GC content (Table 1) fits into the general patern of the GC along the phylogenetic tree (Figure 3).

Dating analysis based on the calibrations known for the primate-associated lice produced an estimate of approx. 6.5 Mya for the split between the N and S lineages of *P. serrata*. This dating places the origin of the N and S lineages considerably deeper than the estimate 1.5 Mya reported previously (Stefka & Hypsa, 2008). Similar to the phylogenetic reconstruction described above, our time estimate obtained here is based on considerably larger data than in the previous study (1049 genes compared to 3). Moreover, the branch lengths of the *P. serrata* lineages shown in Figure 3 are comparable to those of different *Pediculus* species. This estimate and the argument should however be considered with caution. On one hand, the comparison between the two mcmctree runs (regression of the date series; Supplementary figure 2) shows that the analysis reached good convergence and the estimates are not affected by sampling error. On the other hand, however, the estimate falls into a broad confidence interval (95%HPD = 1.8 to 11 Mya). Moreover, the calculation is based on calibration points available only for the primate-associated cluster, presuming similar evolutionary tempo in both louse clusters shown in Figure 3.

Regardless of the exact divergence time, the *P. serrata* lineages N and S seem to represent two distinct species, which differ in an important parameter of their lifestyles, namely host specificity/spectrum (with the S lineage strictly specific to a single host *Apodemus flavicollis*, while N lineage capable to live also on *A. sylvaticus*). Retention of a close morphological similarity over millions of years is not exceptional (Shin & Allmon, 2023). Also, the two *Pediculus* species human louse *P. humanus* and chimpanzee louse *P. schaeffi* are difficult to distinguish despite approx. six millions of years of independent evolution (Reed, Light, Allen, & Kirchman, 2007), although a few morphological features differentiate them (e.g. the width of the thorax). In connection to the *P. serrata* lice, it is pertinent to note that it is extremely difficult (in some cases impossible) to distinguish morphologically also their two host mouse species, estimated to have diverged approximately 4 mya (Michaux & Pasquier, 1974 ex Michaux, Libois, & Filippucci, 2005). Thus, it is possible that the morphological similarity of *P. serrata* hosts and the fact that they share one of the hosts (*A. flavicollis*) conserved morphology of the N and S lineages via stabilizing selection.

### Population demography

To reveal congruence or divergence in coalescence rates for genomic data between the S and N lineages, we used MSMC2 software (Schiffels, Stephan, & Wang, 2020), which estimates effective population size (N_e_) changes during time using Markovian approach, whilst taking into account the connections between linked SNPs. Thus, it reconstructs demographic information not only from coalescence, but considering recombination events as well. Demographic history analyses of both S and N lineages revealed considerable differences in population sizes during the same period of time (Fig. 5). *Polyplax* S showed gradual decline of N_e_ in the past, whereas Polyplax N suffered a more dramatic population collapse with subsequent increase. The lack of possibility to robustly phase *Polyplax* genome did not allow us to relate changes in N_e_ to exact historical events (Schiffels and Durbin, 2014). Moreover, due to the absence of mutation rate estimate in closely related taxa, coalescence times were rescaled using the mutation rate of *Drosophila melanogaster*, which together with the unphased genome reduced the ability to provide absolute timing of demographic events. However, possible errors in timing do not affect interpretation of relative differences in N_e_ over time between the lineages. Life history traits (such as lifespan and fecundity) were proposed to represent the most important drivers of effective population size changes in animals (Romiguier et al., 2014). Given that we compared two closely related lineages with similar life history strategies, except for the difference in host specificities, we expect the shape of the Ne curves to be mostly influenced by varying demographic histories of their hosts. It was shown that the two host species reacted in a different way to the last glaciation period, retreating to different refugia (possibly more fragmented for *A. sylvaticus* than for *A. flavicollis*), and had different re-colonization history (Michaux et al., 2005). In accordance with that, after the glaciation-related decline, the *Polyplax* N lineage could have been able to restore its population size more quickly (when both its hosts recolonized central and northern parts of Europe) compared to the *Polyplax* S, which retained strict specificity to a single host. Correspondingly, our earlier microsatellite-based study (Martinů et al. 2018) showed consistent population genetic diversity differences across several sympatric pairs of *Polyplax* N and S lineage populations, with the S lineage always possessing lower local diversity than the N lineage.

**Figure 5.**
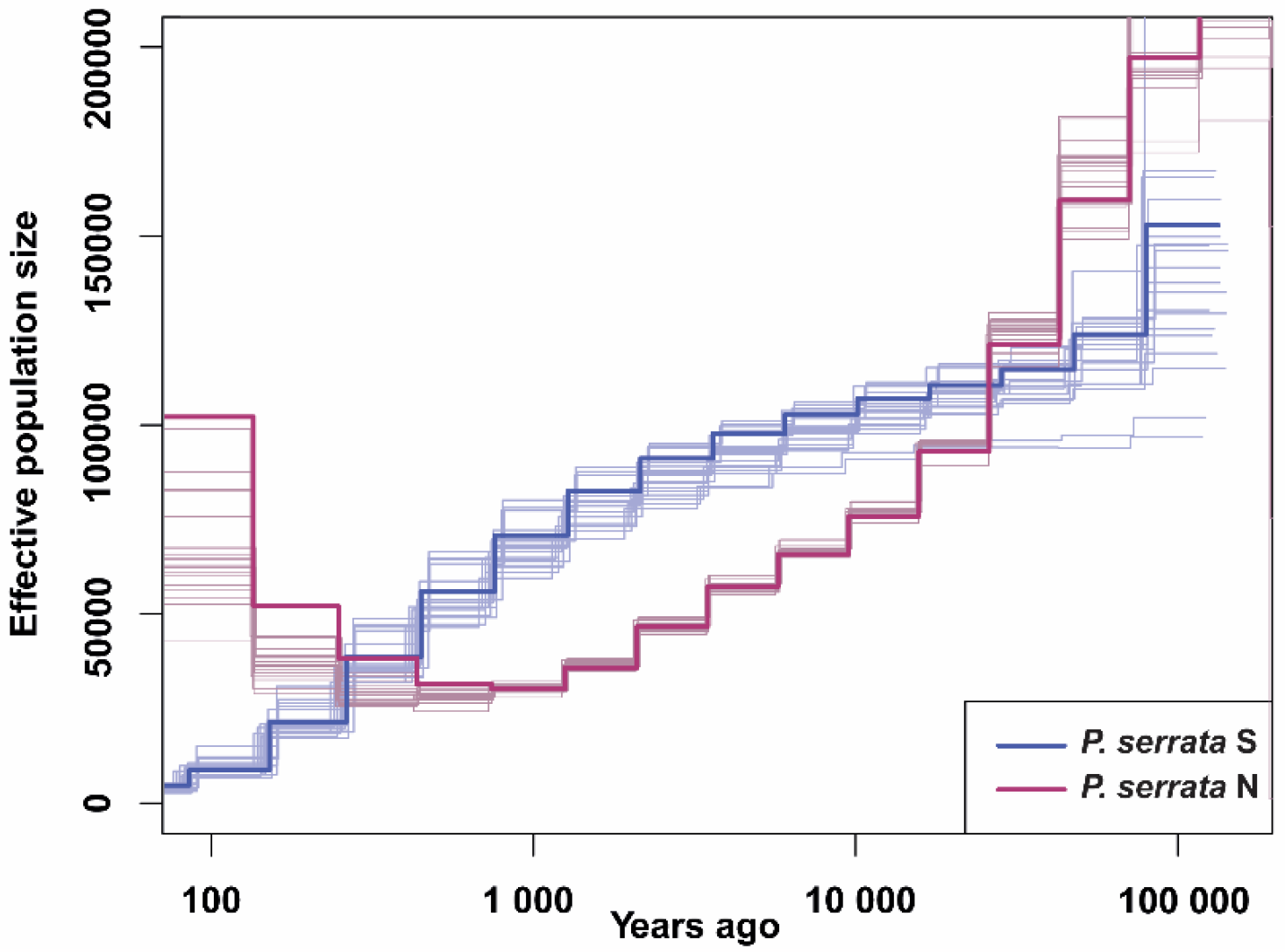
Changes of effective population sizes through time evaluated by MSMC2 for P. serrata S (bold blue) and P. serrata N (bold red) lineages. Boostrap replicates (20 for each lineage) are plotted in lighter lines. Coalescence rates were rescaled using the mutation rate of Drosophila according to Wang et al. (2023) (3.3 × 10–9) and generation time was estimated as 12 generations per year.

## Methods

### Tissue samples and isolation of genetic material

*Polyplax serrata* lice were collected in the northwest of the Czech Republic (CZ) and the Šumava mountains region (CZ) from *Apodemus flavicollis* hosts caught in wooden snap traps. Permission for field studies was provided by the Commitee on the Ethics of Animal Experiments of the University of South Bohemia, by the Ministry of the Environment of the Czech Republic and by the Ministry of the Agriculture of the Czech Republic (Nos. MZP/2017/630/854, 43873/2019-MZE-18134, MZP/2021/630/2459). Lice were gathered from the mouse fur by brushing and stored in 100% ethanol at −20°C. Genomic DNA from individual louse specimens was extracted using the Qiagen QIAamp DNA Micro Kit (Qiagen). Before the next steps, lice were assigned to S and N lineages by sequencing a fragment of the mitochondrial cytochrome oxidase subunit I gene (COI, 379 bp) as in (Martinu et al., 2018).

### Oxford Nanopore and Illumina genomic DNA and RNA sequencing and assembly

One specimen from each lineage (98c_Pro_SE for S lineage, and HR10 for N lineage) with sufficient concentration measured on Qubit 2.0 Fluorometer (Invitrogen) were sequenced using Oxford Nanopore Technology (ONT) on one MinION II flowcell. Preparation of Oxford-Nanopore 1D low input libraries and sequencing were performed at the Roy J. Carver Biotechnology Center (University of Illinois at Urbana-Champaign, USA). Altogether 16.7 Gbp of data were produced for the S lineage and 10.3 Gbp for the N lineage (3.3 and 2.6 million reads of average size of 5 kbp). Datasets were basecalled with Fast-Bonito basecaller (Xu et al., 2021). Raw reads were trimmed with filtlong (version 0.2.0; https://github.com/rrwick/Filtlong) to obtain reads longer than 4,000 bp and with a phred score of 20 and higher. Genome assemblies were reconstructed with flye assembler (version 2.5; (Kolmogorov, Yuan, Lin, & Pevzner, 2019)) using ONT reads with genome size estimation parameter of 110 Mbp. Assemblies were subsequently corrected once with racon (version 1.3.3; https://github.com/isovic/racon) and twice with medaka (version 0.6.5; https://nanoporetech.github.io/medaka/) using the ONT reads. The last polishing step was performed again with racon, this time using Illumina reads obtained from the same specimen as ONT reads. Illumina reads from S and N samples were sequenced on a Novaseq lane as part of a different study and details are described in (Martinu et al., 2020). Completeness of the assemblies were checked with BUSCO using Arthropoda gene set (version 3; (Waterhouse et al., 2018).

For RNA sequencing of the S lineage, 34 lice of different life stages were gathered from one specimen of the field mouse *Apodemus flavicollis* caught in the proximity of a game preserve Flaje (CZ). Lice were preserved in RNAlater (Thermo Fischer Scientific) and isolated with phenol-chloroform extraction in the laboratory. Preparation of the cDNA library and sequencing of the 150bp long PE reads on Illumina NovaSeq 6000 sequencer were performed by Novogene (United Kingdom). Adapters and low-quality reads were removed using Trimmomatic v0.39 (Bolger, Lohse, & Usadel, 2014). Obtained reads were then checked for quality with FastQC (Andrews, 2010) and assembled with Trinity v2.15.1 with default settings (Grabherr et al., 2011). RNA data for N lineage were not obtained due to lack of sufficient amount of input material.

Since the assemblies combining Illumina and Oxford nanopore reads contained genomes of the symbiont *Legionella polyplacis* fragmented into several contigs, we used (based on our previous experience) Spades 3.10 (Bankevich et al., 2012) to assemble complete symbionts’ genomes from the Illumina short reads. The reads were trimmed by Trimmomatic (Bolger et al., 2014) with default parameters. Spades was run with the option --meta. In the resulting assemblies, the contigs representing complete *L. polyplacis* genomes were identified based on their lengths and ORFs distribution (i.e., densely arranged ORFs as typical for prokaryotes), and were annotated in Prokka (Seemann, 2014).

### Gene prediction, annotation, and functional comparative analysis

To compare our genomes with the other three available phthirapterans, we reannotated and reanalysed all genomes, rather than using the published data for the comparison. This ensures a more consistent approach. The results we obtained from these reannotations differ in some aspects from those published previously, particularly in the numbers of genes identified in various categories and families. Considering the complexity of eukaryotic genomes, the level of uncertainty during the annotation process, and the differences in annotation approaches/programs, such inter-study differences are to be expected. However, they also provide warning that the results of the annotation step and idenfication of gene functions must be taken with caution (Salzberg, 2019; Scalzitti, Jeannin-Girardon, Collet, Poch, & Thompson, 2020).

To perform genomic comparison of *P. serrata* with other Phthiraptera, we included into our analyzes the three previously published genomes, i.e., *Pediculus humanus, Brueelia nebulosa*, and *Columbicola columbae*. While there are three other phthirapteran genomes deposited in the NCBI GenBank, we did not include them due to their lower quality. To compare several specific genes/functions of the studied lice with other blood feeding insects, we also included genomes of *Aedes aegypti, Cimex lectularius, Glossina morsitans*, and *Rhodnius prolixus* (See Supplementary Table S1, Datasheet S8 for the NCBI GenBank accession numbers and transcriptome references of the genomes). As a preparatory step for the gene prediction, we identified repeat contents in our de novo assemblies of *P. serrata* utilizing RepeatModeler v2.0.3 (Flynn et al., 2020), which was followed by soft masking complex repeats using RepeatMasker v4.1.2-p1 (Tarailo-Graovac & Chen, 2009). To maintain a methodologically consistent approach to the down-stream analysis, we applied the same prediction and annotation process to all of the included genomes. This procedure ensured methodological uniformity, and minimized variability that might arise from using different tools, databases, or varying versions of the databases in the previous studies. Funannotate v1.18.14 (https://github.com/nextgenusfs/funannotate) was employed to perform gene prediction and functional annotation on the analyzed genomes. Briefly, *ab initio* gene prediction was performed by “funannotate predict” command that employed Augustus v3.5.0 (Hoff & Stanke, 2019), glimmerHmm v3.0.4 (Majoros, Pertea, & Salzberg, 2004), snap v2013_11_29 (Kolesov, Mewes, & Frishman, 2001) and GeneMark v4.71 (Lukashin & Borodovsky, 1998) gene predictors. Transcript assemblies were used as transcript evidence to enhance gene prediction (with exception of *Brueelia nebulosa* and *Glossina morsitans* for which no transcriptomic evidence is available in public databases). The derived gene models underwent annotation via the “funannotate annotate” command, which invokes InterProScan v5.60-92.0 (Zdobnov & Apweiler, 2001), eggNOG-mapper v2.1.10 (Cantalapiedra, Hernández-Plaza, Letunic, Bork, & Huerta-Cepas, 2021), SignalP v5.0b (Almagro Armenteros et al., 2019) and Phobius v1.0.1 (Kall, Krogh, & Sonnhammer, 2004) tools for gene annotation. Comparative analysis was performed using the “funannotate compare” function of Funannotate v1.8.14 (https://github.com/nextgenusfs/funannotate).

### Comparative functional genomic analysis of Anoplura and chewing lice

To elucidate shared and unique protein families and domains across the compared Anoplura (*P. serrata* S and N lineage and *P. humanus*) and chewing lice (*C. columbae* and *B. nebulosa*) genomes, Venn diagrams were generated based on functional comparison outputs (Supplementary Table S1, Datasheets S3-S5, Datasheet S7) from different protein annotation databases. This included Pfam and InterPro (Paysan-Lafosse et al., 2023) databases for annotated protein families, domains and conserved sites, the carbohydrate-active enzyme (CAZy) database for classifications of carbohydrate active enzymes (Drula et al., 2022) and MEROPS database for peptidases and peptidase inhibitors . In addition, a principal coordinate analysis (PCoA) was conducted on comparison output from the Pfam and InterPro databases (Supplementary Table S1, Datasheets S3-S4), which utilized Bray-Curtis dissimilarity metric in vegan package v2.6-4 (Oksanen, 2011) within R environment (R Core Team, 2013).

### Metabolic reconstructions of host-symbiont complementarity

To evaluate and compare possible complementarity of the louse hosts and their obligate symbionts in production of amino acids and B vitamins (the compounds typically considered in the insect-bacteria symbiosis), we analyzed metabolic capacities using the KEGG database (Kanehisa, Sato, Kawashima, Furumichi, & Tanabe, 2016) . For the genomes of both *P. serrata* lineages and their *L. polyplacis* symbionts, we assigned K numbers (KEGG orthology identifiers) to all annotated proteins in their genomes by the web-based program BlastKoala (Kanehisa, Sato, & Morishima, 2016), and mapped these numbers on the biosynthetic pathways using the KEGG mapper tool. For *P. humanus* and its symbiont *R. pediculicola*, we used the pathway maps already available in the KEGG database.

### Phylogeny and evolutionary dating of the P. serrata *lineages*

To reconstruct phylogenetic position of the two *P. serrata* linages within Anoplura, we build a matrix of 15 Anoplura species and two sequences of *Haematomyzus elephantis* as outgroups. Using a locally generated pipeline (refer to the Data availability section), we extracted orthologs from the alignment published by de Moya et al. (2021) in their phylogenomic analysis of Psocodea. (Downloaded from https://datadryad.org/stash/dataset/doi:10.5061/dryad.c59zw3r50. Since we focused on Anoplura, we selected for the following analysis only the 13 anopluran species and the outgroup. These data were extended with the genes extracted from the *P. serrata* genomes. To obtain sets of single copy orthologs, we translated all sequences into amino acids using the Emboss v6.6.0.0 (W. Li et al., 2015) transeq function, and searched the orthologs by Orthofinder v2.5.5 (Emms & Kelly, 2015). For the total of the 1049 identified single copy orthologs, we made alignment of their nucleotide forms using Mafft (Katoh, Misawa, Kuma, & Miyata, 2002) implemented in Geneious (Kearse et al., 2012). The alignments were concatenated and a matrix was built from all second codon positions, resulting in a matrix of 536,147 positions. The tree was inferred using iqtree2 v. 2.2.0 (Minh et al., 2020) by maximum likelihood under the GTR+F+I+I+R3 model chosen by program based on BIC, with 1,000 ultrafast bootstrap replicates. The resulting topology was further used as a constraint for the dating analysis in the mcmctree program (Puttick, 2019). Calibration for two nodes within Anoplura was adopted from the (de Moya et al., 2021) analysis, namely *Pedicinus* + (*Phthirus* + *Pediculus*) (20-25 Ma), and *P. schaeffi* + *P. humanus* (5–7 Ma). To derive the estimates for the other nodes, we ran the mcmctree analysis with approximate likelihood calculation and the mcmc process set to 500,000 samples, burnin 50,000 and sample frequency 10. Convergency of the mcmc chains was checked by comparing convergence between the dating sets from both chains, as recommended in the mcmctree manual.

### Synteny analysis

For comparative analysis of *Polyplax* S and N lineages, contigs longer than 0.7 Mbp were chosen. The level of synteny of the 18 longest scaffolds of *Polyplax* S and 21 of *Polyplax* N was analyzed using McScanX (YP Wang et al., 2012), method considering orthologous genes as anchors. Collinearity for the orthologue synteny blocks on contigs was evaluated using default settings. SynVisio program (Bandi & Gutwin, 2020) was used for detailed visualization of the McScanX outputs in regions where structural rearrangements occurred.

### Population demography

To compare population demography of the two *P. serrata* lineages, we prepared genomic DNA libraries for individual louse samples with insert size of 450 bp. The libraries were sequenced by paired-end Illumina process on NovaSeq6000, yielding approximately 59.5 million paired-end reads per sample. Preparation of libraries as well as sequencing were provided by the W. M. Keck Center (University of Illinois, Urbana, IL, USA). For the following analysis, we selected eight samples for each lineage (the S lineage data were also part of the previously published study; (Martinu et al., 2020). The demographic history of both lineages was evaluated from the whole genome sequences by Multiple Sequentially Markovian Coalescent (MSMC2) software (Schiffels et al., 2020). The data were adaptor and size filtered with bbtools (https://jgi.doe.gov/data-andtools/bbtools/) and reads were mapped against *Polyplax* S genome using bowtie2 (Langmead & Salzberg, 2012). Duplicated reads were removed with PICARD (http://broadinstitute.github.io/picard/) and SNP calling was performed using GATK Genome Analysis Toolkit following the “Best Practices” guide from the Broad Institute (Van der Auwera et al., 2013). Datasets of *Polyplax* S and N were separately filtered for quality with GATK, then minor allele frequencies (MAF) equaled to 0.05 were removed in PLINK 1.9 (https://www.cog-genomics.org/plink/1.9/). Data were then converted according to MSMC2 instructions using SAMtools v.1.8 (H. Li et al., 2009), BCFtools v.1.8 (Danecek et al., 2021) and scripts available at (msmc-tools/msmc-tutorial/guide.md at masterstschiff/msmc-tools·GitHub). MSMC2 analyses assessed coalescence rates between haplotypes within the S and N lineages as well as 20 bootstrap replicates based on default values except for time segment patterning parameter (-p). To avoid overfitting, the default 32 time segments were lowered to 18 (-p 1*2+15*1+1*2), due to the small size of the *Polyplax* genome (139 Mbp). Results were plotted in R Studio, where they were scaled based on the mutation rate of *Drosophila* (3.3 x 10^−9^) (YG Wang et al., 2023)(Wang et (Van der Auwera et al., 2013)al., 2023) and generation time of 12 generations per year.

## Data availability

The annotated genomes of *P. serrata* S and N lineages, the re-sequenced samples for the demography reconstruction, and the genomes of *Legionella polyplacis* were deposited on GenBank database under BioProject PRJNA1018720, with GenBank accession numbers JAWJWF000000000, JAWJWE000000000 and CP135136, CP135137, respectively. Datasets of *P. serrata* S and N lineages and other compared genomes are available at Zenodo (https://zenodo.org/records/8381480) under doi:10.5281/zenodo.8381480. This includes fasta format files of genomes, transcriptome and proteome and annotation tables obtained from gene prediction and annotation workflow. In addition, Zenodo-deposited datasets include gbk format files for all compared genomes and identified repeat families in fasta format files for *P. serrata* S and N lineages. Helper python, bash and R scripts employed in this study are available at https://github.com/hassantarabai/MS-Pserrata-S-N-2023.

## Acknowledgements

We thank our colleagues Masoud Nazarizadeh, Jakub Vlček, and Milena Nováková for their advice on bioinformatic procedures. Access to computing and storage facilities owned by parties and projects contributing to the National Grid Infrastructure MetaCentrum provided under the program “Projects of Large Research, Development, and Innovations Infrastructures” (CESNET LM2015042) is greatly appreciated. This work was supported by the Grant Agency of the Czech Republic (grant number 21-02532S to V.H.).

## Supplementary figures

**Supplementary figure 1.**
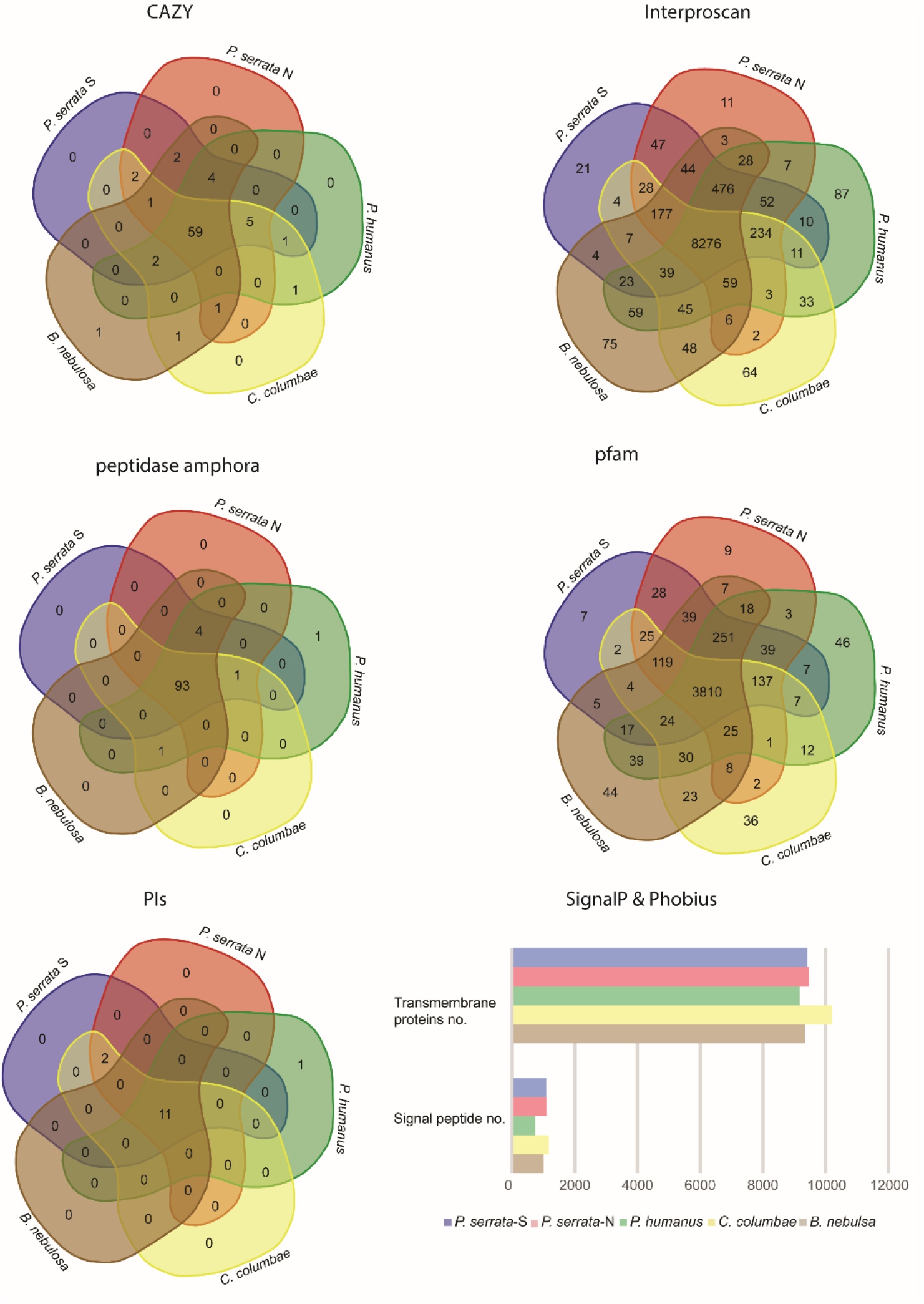
Comparison of genome contents of the five lice (Phthiraptera). For the databases CAZy, InterProscan, MEROPS, Pfam, the numbers represent unique IDs identified in the genomes. For SignalP & Phobius the plot shows total numbers of the transmembrane proteins and signal peptides identified by the databases.

**Supplementary figure 2.**
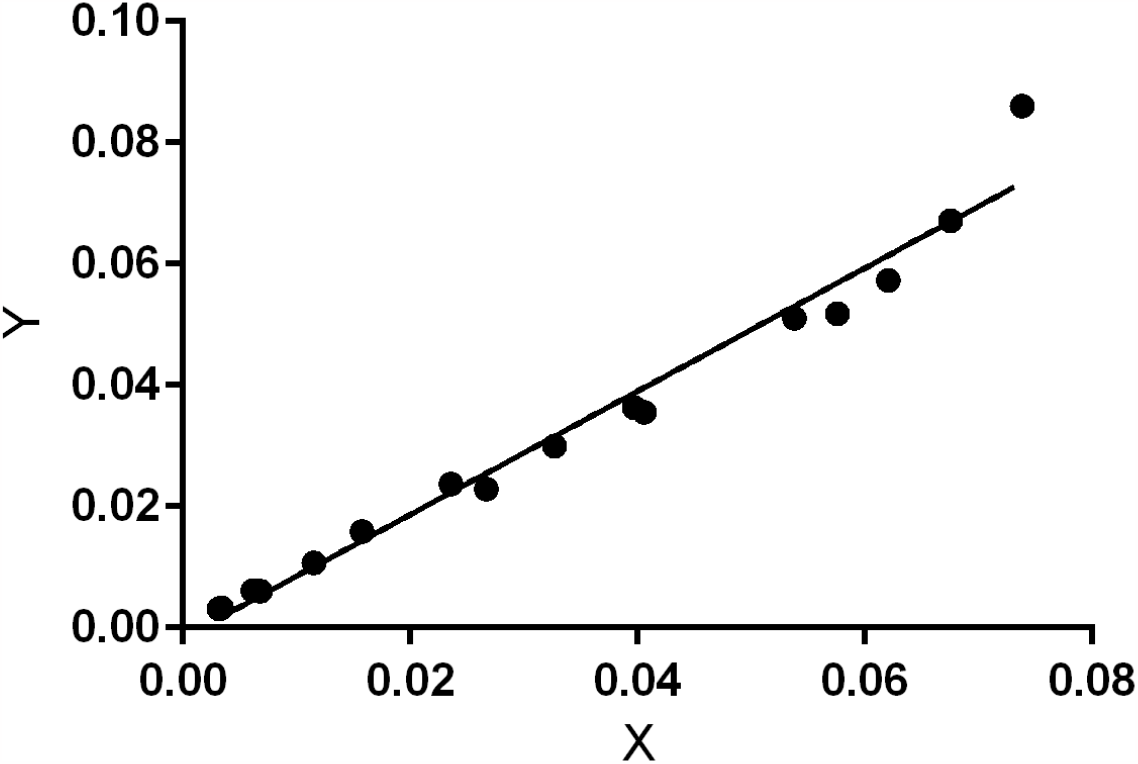
Convergence of two independent runs of mcmctree analysis demonstrated on plot of the posterior times produced by GraphPad (https://www.graphpad.com/quickcalcs/linear1/). The correlation coeficient = 0.9992

